# Heterogeneous responsiveness to environmental stimuli^*^

**DOI:** 10.1101/2023.01.26.525694

**Authors:** Jerome Cavailles, Christoph Kuzmics, Martin Grube

**Affiliations:** Institute of Biology, University of Graz, Graz, Austria; Institute of Economics, University of Graz, Graz, Austria

**Keywords:** equilibrium, ESS, behavioral ecology, individual differences

## Abstract

Individuals of a species cope with environmental variability through behavioral adjustments driven by individuals’ responsiveness to environmental stimuli. Three key empirical observations have been made for many animal species: The *coexistence* of different degrees of responsiveness within one species; the *consistency* of an individual’s degree of responsiveness across time; and the *correlation* of an individual’s degree of responsiveness across contexts. Taking up key elements of existing approaches, we provide one unifying explanation for all three observations, by identifying a unique evolutionarily stable strategy of an appropriately defined game within a stochastic environment that has all three features. Coexistence is explained by a form of negative frequency dependence. Consistency and correlation is explained through potentially small, individual, differences of states animals have and the resulting differential advantages they can get from it. Our results allow us to identify a variety of testable implications.

## 1 Introduction

Individuals generally cope with frequent changes in their environment [Candolin et al., 2023] by means of behavioral adjustments, see e.g., Coulson et al. [2017], Sih et al. [2012], Wong and Candolin [2015], Sih [2013]. To be able to adjust their behavior to changes in the environment, individuals need to be responsive to environmental stimuli, observable indicators of these changes.^1^

It is well documented that individuals differ in the degree of responsiveness to external stimuli, a phenomenon sometimes referred to as behavioral plasticity, see e.g., Mitchell and Houslay [2021], Laskowski et al. [2022]. More generally, *phenotypic plasticity* – which use the alterations in an organism’s behavior, morphology, and physiology as a response to its specific environmental conditions – have been extensively studied. Phenotypic plasticity was once undervalued and largely overlooked as a mechanism and concept of evolution. However, a shift in perspective is occurring due to recent theoretical and empirical studies. These studies highlight the importance of plasticity in fostering novelty and diversity across morphological, physiological, behavioral, and life history traits. For recent overviews, see Pfennig [2021] and Sommer [2020]. The concept of reaction norms originates from Woltereck [1909].

This difference in environmental responsiveness constitutes the main characteristic of personalities, and personalities have been observed in more than 100 species, see e.g., the survey by Nettle and Penke [2010]. In a broader sense, animal personality can be related to plasticity [Dingemanse et al., 2010]. Three key observations have been made for many animal species, as, for instance, highlighted by Wolf et al. [2007], see also Bell et al. [2009]: The *coexistence* of different degrees of responsiveness within one species; the *consistency* of an individual’s degree of responsiveness across time; and the consistency, often referred to as *correlation*, of an individual’s degree of responsiveness across contexts.^2^

A few theoretical approaches explain one or more of these three observations. The theory of biological sensitivity to context, as in Pluess [2015], Boyce [2016], and Ellis and Boyce [2008] explains the coexistence of different degrees of environmental responsiveness with differences in individuals’ experiences in their early development. The theory of differential sensitivity, as in Ellis et al. [2011], interprets the difference of behaviour as a way to hedge future generations against the uncertainty in the environment, recently formalised by Bergstrom [2014] and Frankenhuis et al. [2016].^3^ A third approach is built around the idea of “negative frequency dependence:” The more individuals are responsive to environmental stimuli the less the benefits of being responsive. Negative frequency dependence is a cornerstone for explanations of the coexistence of different degrees of environmental responsiveness in the seminal models of Wolf et al. [2008], Wolf et al. [2011], and Wolf and McNamara [2012]. See Dingemanse and Wolf [2010] for a review of earlier models. While negative frequency dependence is able to explain coexistence, consistency and correlation are often explained by an individual’s state (e.g., morphology, phenotype, size, etc.), as in Wolf and Weissing [2010] and Dingemanse and Wolf [2010].^4^ However, the meta analysis by Niemelä and Dingemanse [2018] shows a weak link between state and personalities (individuals’ state can only explain between 3 and 8% of the personality differences).

We study individual responsiveness to environmental stimuli for the specific problem of foraging from multiple food sources, which allows us to additionally build on the existing game-theoretic literature on the *ideal free distribution* of Fretwell and Lucas [1969], see also Křivan et al. [2008]: individuals allocate themselves proportionally to the amount of food available at each food source. For a more general framework about foraging theory see Stephens and Krebs [1986], Pyke [2019]). We enrich these game-theoretic models by embedding them in a stochastic environment, which allows us to incorporate the salient features of the three approaches mentioned above. We do this in a few steps of varying complexity.

The basic model (provided in Section 2) is sketched in Figure 1. To illustrate the model we use fish-feeding birds as an example (inspired from observations of Wakefield et al. [2015], McHuron et al. [2018], Harris et al. [2020], Patrick et al. [2014]), with the simplifying condition that they do not show any social behavior (such as flocking or swarming). Individuals have to choose to forage from one of two food sources, one providing a fixed amount of food, the other a random amount of food. Individuals can, at some small cognitive cost, learn about the state of the food availability at the random source. All individuals who go to the same food source are assumed to share the available food there equally. In the basic model the random food source can only have two possible levels of food availability.

**Figure 1:**
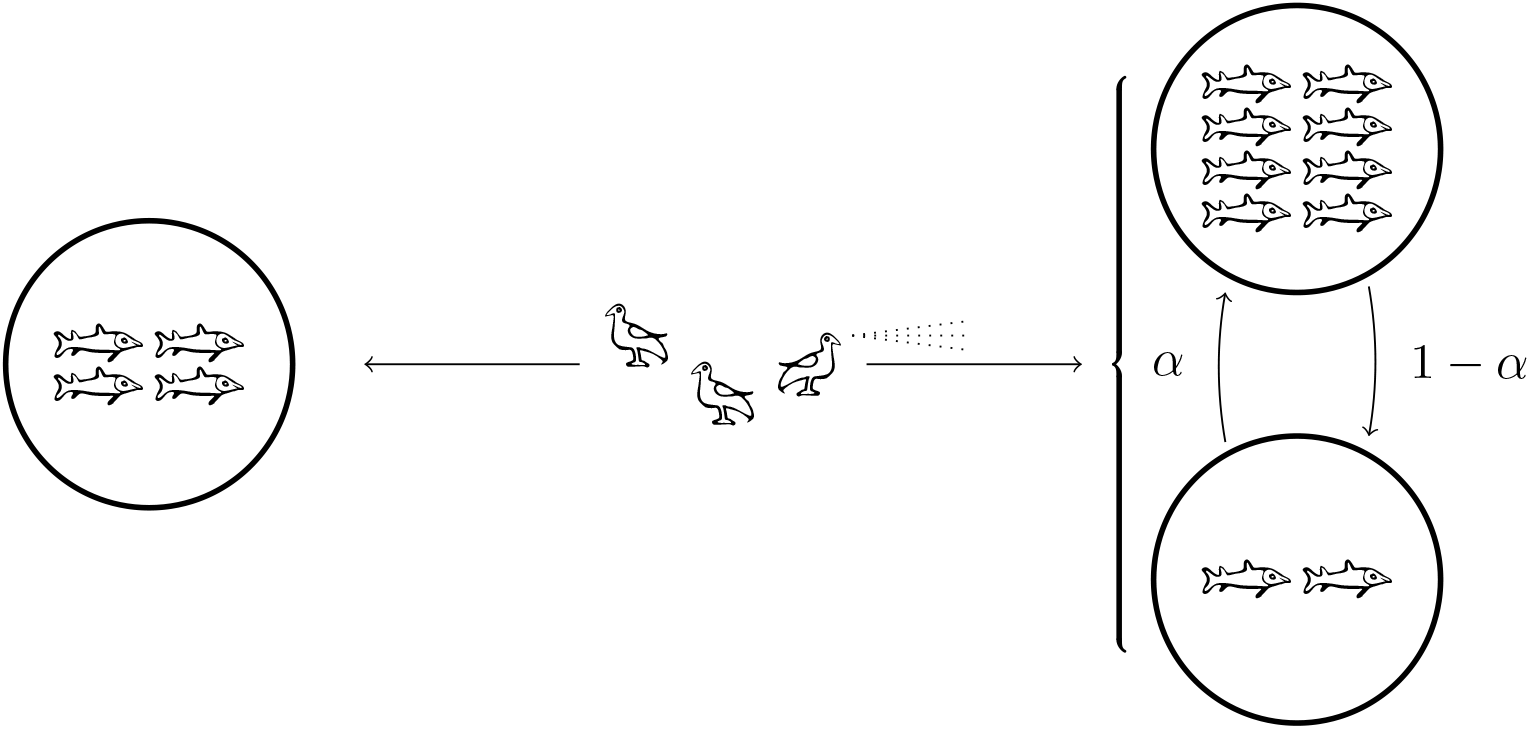
A graphical sketch of the basic model. The birds are the players, who choose which food source to go to: fixed food source A on the left or stochastic food source B on the right, with *α* the probability of high food availability. The bird’s scanning indicates that players can make their decisions based on observing the food availability at the stochastic source. In this concrete example, at the equilibrium, one bird should always go to the left, one bird should always go to the right, and one bird should be responsive to change. They will all eat two fishes when the stochastic source is low, and four fishes when the stochastic source is high.

Our first main result is that the game always has a unique symmetric Nash equilibrium strategy, Nash [1950], and this strategy exhibits coexistence of responsive and non-responsive individuals. This equilibrium is also evolutionarily stable, an ESS, in the sense of Palm [1984], who extended the definition of an ESS by Maynard Smith and Price [1973] to symmetric *n*-player games, see also Broom et al. [1997]. Moreover, the games are *stable* in the sense of Hofbauer and Sandholm [2009], which implies that the unique symmetric Nash equilibrium is also asymptotically stable under most plausible behavioral adjustment dynamics, including the prominent replicator dynamics of Taylor and Jonker [1978].

The equilibrium is in mixed strategies and individuals are indifferent between being responsive and not being responsive. This raises concerns about the consistency of their behavior (across time and contexts). We then introduce small perturbations to the model by allowing slight differences in individual-specific food preferences and in the individuals’ cost of being responsive. This is the idea of Harsanyi [1973] *purification* and the idea of *threshold decisions* of McNamara and Houston [2005]: The original mixed strategy equilibrium (or ESS) corresponds in the perturbed game to an equilibrium in which every individual has a strict preference to either be or be not responsive. To the extent that these idiosyncratic payoff perturbations are constant over time and across contexts we now also get *consistency* and *correlation* in addition to *coexistence*.

We also provide an analytic expression for the equilibrium pure strategy frequencies as a function of the parameters of the problem. This allows us to derive additional testable predictions of our model. Acknowledging that our model is a highly simplified account of reality, some of these predictions may yet hold beyond the narrower confines of our model. In particular, and perhaps most striking: the unique equilibrium does not change with the stochasticity of the environment, at least when the cost of cognition is negligible. This implies that changes to the stochastic process of the environment may not push behavior out of equilibrium. Individuals’ strategies are already sufficiently complex to allow essentially immediate and automatic adaptation to such changes. Of course, this does not imply that individuals are not affected if, for instance, high food availability becomes rarer. Only their strategy is unaffected, not necessarily the amount of food they can consume.

Another testable prediction can be derived from an extension of the basic model, in which individuals only receive (private) noisy information about the state of food availability. At least for small amounts of noise, the equilibrium responsiveness increases when the noise increases. More noise, in some sense, forces individuals to overreact to environmental stimuli. A final testable prediction, again derived from an extension of our basic model, is that for more general distributions of food availability, one would expect to see a continuum of degrees of responsiveness to environmental stimuli. See e.g., Lionetti et al. [2018] for empirical support for this finding.

The rest of the paper proceeds as follows. Section 2 provides the basic game-theoretic model. Section 3 derives the unique symmetric Nash equilibrium of the relevant games. This equilibrium exhibits “coexistence” of different degrees of environmental responsiveness. Section 4 provides a further discussion of the implications of equilibrium behavior. Section 5 then extends the model slightly to accommodate both “consistency” and “correlation” of individuals’ degree of responsiveness, across time and contexts, respectively. Section 6 considers the case of noisy information, and Section 7 provides a generalisation of the basic model to general distributions of food availability at the stochastic source. Section 8 finally concludes with a further discussion of the results and further related literature. The Appendix provides some of the more technical arguments behind some of the results in the paper.

## 2 Methods

In this section, we present the simplest possible model of interest for our problem. Each of *n < ∞* individuals can go to one of two food sources *A* or *B*. Food source *A* has a fixed amount of food normalized to *n* units of nutrition to facilitate an easier comparison when we vary the number of individuals *n*. Food source *B* has a stochastic amount of food *n · X*, with *X* drawn from a Bernoulli distribution with *X* = *η* with probability *α* and *X* = *λ* with probability 1 − *α* with *λ* denoting a low and *η* a high food-availability at the stochastic food source *B*. We assume that 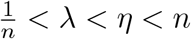. If the number of individuals *n* is large, the assumption represents barely a restriction. If *n* is not very large, the assumption represents the most interesting case. Suppose, for instance, that both *λ*, 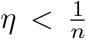. This would imply that, the food-availability at the stochastic source *B* is always less than 1, while the food-availability at source *A* is *n*. Thus, as we are assuming equal food-sharing (see below), no individual would ever benefit from going to the stochastic food source. Even if all individuals go to food-source *A* an individual who switches to the stochastic source *B* gets a food-share of 1 only. Suppose, as another extreme, that both *λ, η > n*. Then the food-availability at the stochastic source *B* always exceeds *n*^2^ and, therefore, no individual would ever benefit from going to the fixed source *A*. They get more than *n* units of nutrition at food source *B* and only at most *n* at food source *A*.

Before making their choice of food source, individuals can, in principle, inform themselves about the state of food source *B*; individuals can choose to learn whether *X* = *λ* or *X* = *η*. If uninformed they can then choose to go to food source *A* or *B*. If informed they can react to the information in one of two ways. ^5^ They can be, what we term, *responsive* by going to *A* when *X* = *λ* and going to *B* when *X* = *η*, or, what we term *counter-responsive* by doing the opposite. We denote the set of strategies by *S* = *{A, B, R, C}*, for always going to food source *A*, always to *B*, being responsive, and being counter-responsive, respectively.

**Table 1:** Overview of key notation.

| Symbol | Description | Range |
| --- | --- | --- |
| parameters |  |  |
| $n$ | number of individuals | $\mathbf{N}$ |
| $\lambda$ | lower amount of food at the variable food source | $\mathbf{R}^{*+}$ |
| $\eta$ | higher amount of food at the variable food source | $\mathbf{R}^{*+}$ |
| $\alpha$ | probability to have a high amount of food at the source $B$ | $[0, 1]$ |
| $c$ | cost for being responsive (or counter-responsive) | $\mathbf{R}^+$ |
| $X$ | Stochastic variable of the amount of food at the source $B$ | $\{\lambda, \eta\}$ |
| strategies |  |  |
| $S$ | Set of strategies | |
| $A$ | Strategy *always go to food source A* | |
| $B$ | Strategy *always go to food source B* | |
| $R$ | Strategy *being responsive* | |
| $C$ | Strategy *being counter-responsive* | |

Let Δ(*S*) denote the set of all mixed strategies, that is the set of all probability distributions over *S*. There are two ways to interpret a mixed strategy *σ* ∈ Δ(*S*), both go back at least to Maynard Smith and Price [1973] who speak both of the *probability* and the *frequency* of certain actions.

One can think of a mixed strategy as a way an individual randomizes over pure strategies. These pure strategies are then played with the probability specified by *σ*. But one can also think of a mixed strategy as resulting from a random draw of a large (essentially infinite) population of individuals with given proportions of them playing the various pure strategies. These pure strategies are then present in the population with frequencies given by *σ*. While mathematically equivalent, this distinction matters for how our results are able to explain both the consistency and correlation of behaviour, which we investigate in the respective section below.

We assume throughout the paper that all individuals, who go to the same food source, share the available food at this source equitably. The payoff to an individual then only depends on the number of other individuals, *k*, that go to food source *A* (which implies that *n* − 1 − *k* others go to food source *B*). The payoff to an individual who goes to food source *A* is given by 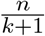; the payoff to an individual who goes to *B* is given by 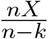.

Given that individuals can choose mixed strategies, we need to compute individuals’ expected payoffs. To do so, consider an arbitrary individual who is facing that all other *n*− 1 individuals choose a given (mixed) strategy *σ* ∈ Δ(*S*) with *σ*(*s*) the probability that pure strategy *s* is chosen.

Denote by *N*_*s*_ the random variable that is the number of opponents who end up choosing pure strategy *s* ∈ *S* (given the probability of choosing *s* is *σ*(*s*)). The tuple (*N*_*A*_, *N*_*B*_, *N*_*R*_, *N*_*C*_) is then multinomially distributed with the probability vector (*σ*(*A*), *σ*(*B*), *σ*(*R*), *σ*(*C*)).

Given *σ*, let *R*_*Aλ*_ denote the random variable that is the food share available at food source *A* in state *λ*. Let food shares *R*_*Aη*_, *R*_*Bλ*_, *R*_*Bη*_, be defined analogously. These are given by

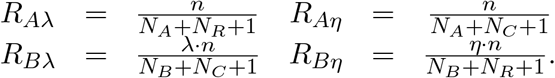

We can, then, express an individual’s expected payoffs from choosing pure strategy *s* ∈ *S*, when all others use mixed strategy *σ*, as follows.

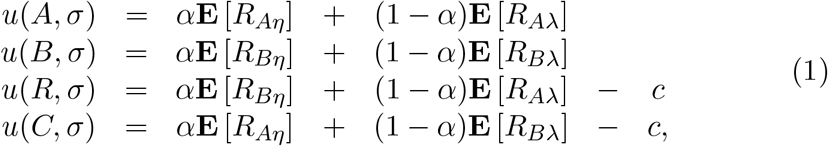

where **E** denotes the expectation with respect to the randomness created by mixed strategy *σ*.^6^ We extend an individual’s payoff function to mixed strategies by taking expectations:

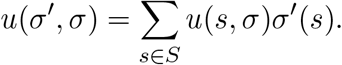

Strategy *σ* ∈ Δ(*S*) is a *symmetric equilibrium* [Nash, 1950] if *u*(*σ, σ*) ≥ *u*(*σ*^*′*^, *σ*) for all *σ*^*′*^ ∈ Δ(*S*).

## 3 Equilibrium

We first show that pure strategies *A, B*, and *R*, cannot be equilibria for any *c >* 0 and for any *n* given the assumption that 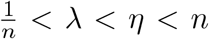 . Suppose everyone plays *A*. Then everyone gets a payoff of 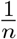 . An individual deviating to *A* would at least get *λ* and since 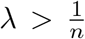 this is a profitable deviation.

Suppose everyone plays *B*. Then everyone gets at most *η < n* (if *X* = *η* and less if *X* = *λ*). Deviating to *B* would yield a payoff of *n*. Suppose, finally, that everyone plays *R*. Then everyone gets a payoff of either 1 (if *X* = *λ* and everyone goes to food source *A*) or *η* (if *X* = *η* and everyone goes to food source *B*), minus *c*, and then a deviation to *B* gives a higher payoff of *λn >* 1 when *X* = *λ* (this individual is alone at food source *B*) and a payoff of *n > η* if *X* = *η* (this individual is alone at food source *A*), and thus, a strictly higher expected payoff.

Next we show that there cannot be a symmetric Nash equilibrium that attaches non-zero probability to both pure strategies *R* and *C*. If this were the case, then necessarily *u*(*C, σ*) = *u*(*R, σ*) and, by the fact that *u*(*A, σ*) + *u*(*B, σ*) = *u*(*R, σ*) + *u*(*C, σ*) − 2*c* for all *σ* ∈ Δ(*S*), either pure strategy *A* or *B* (or both) provide a strictly higher payoff than both *R* and *C*.

There cannot be a symmetric Nash equilibrium in which strategy *C* is used with non-zero probability. To see this, we show that whenever *C* is used *R* must be used with non-zero probability. As we just showed that there cannot be an equilibrium with both *C* and *R*, this then proves that there is no equilibrium in which *C* is used.

We prove this by contradiction. Suppose, that *σ*(*R*) = 0 and *σ*(*C*) *>* 0 for an equilibrium *σ* ∈ Δ(*S*). We will show that, in this case, either strategy *A* or *B* will provide a strictly better payoff than *C* does. Noting that *N*_*R*_ = 0, we can write the expected payoffs as

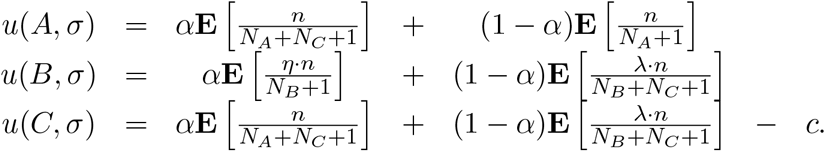

As *σ* is an equilibrium we must have *u*(*A, σ*), *u*(*B, σ*) ≤ *u*(*C, σ*). This implies that

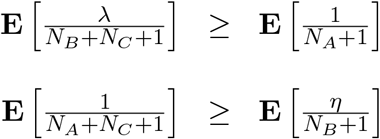

But, since we have E[N_C_] > 0, we have

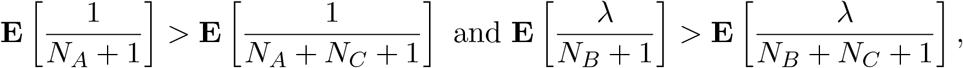

which, in combination with the previous two inequalities yields

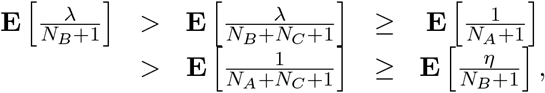

and, thus,

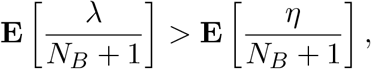

which, finally, contradicts our assumption that *η > λ*.

Next we show that, given the assumption that 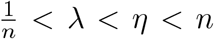, there cannot be any mixed strategy equilibrium with only two pure strategies in its support. We have already shown that there cannot be an equilibrium with *C* in its support. Suppose there is an equilibrium *σ* that has a support of *{A, R}*. Then *u*(*A, σ*) = *u*(*R, σ*). However, *u*(*B, σ*) *> u*(*R, σ*), as *B* provides the same payoff as *R* when *X* = *η* and obtains *λn* when *X* = *λ* (as everyone else is at source *A*), while *R* obtains 1 *< λn*. Suppose there is an equilibrium *σ* that has a support of *{B, R}*. Then *u*(*B, σ*) = *u*(*R, σ*). However, *u*(*A, σ*) *> u*(*R, σ*), as *A* provides the same payoff as *R* when *X* = *λ* and obtains *n* when *X* = *η* (as everyone else is at source *B*), while *R* obtains *η < n*.

Suppose there is an equilibrium *σ* that has a support of *{A, B}*. Then *u*(*A, σ*) = *u*(*B, σ*). As *η > λ*, then for sufficiently small costs *c, R* provides a higher payoff than both *A* and *B*. In expectation both food sources *A* and *B* have the same food share, but *R* provides a higher payoff by going to *B* if and only if *X* = *η*.

This, finally, leaves as the only possible equilibria those with support given by *{A, B, R}*. Let *r*_*yx*_ = **E**[*R*_*yx*_] denote the expected food share at source *y* ∈ *{A, B}* in state *x* ∈ *{λ, η}*. Consider *r*_*Aλ*_. All other expected food shares are calculated analogously. We can write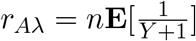, where *Y* = *N*_*A*_ + *N*_*R*_, and *Y*, thus, binomially distributed with *n* − 1 tries and success probability *p* = *σ*(*A*) + *σ*(*R*). The first negative moment of such a binomial distribution, 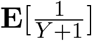, is provided in [Chao and Strawderman, 1972, Equation 3.4] and given by 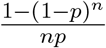.

We, thus, obtain

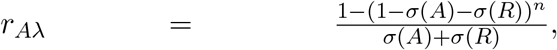

and analogously

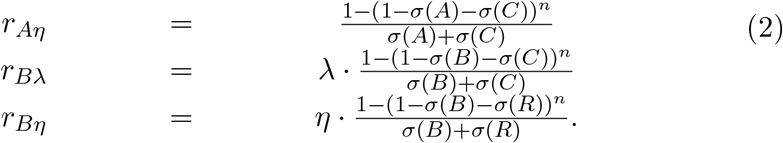

We need to show that there is a unique equilibrium with support *{A, B, R}*. To be indifferent between *A* and *R* and *B* and *R* requires that

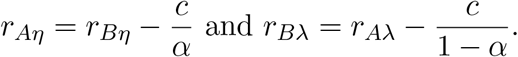

Plugging in the above expressions, and noting that *σ*(*C*) = 0, these two equations lead to

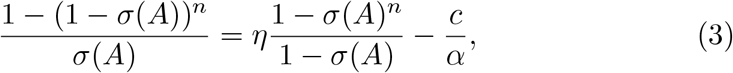

and

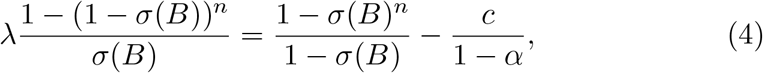

respectively.

In general, we cannot provide an analytic solution to this system of equations. If, however, we consider the limit of the number of individuals going to infinity, *n* → *∞*, the system of equations simplifies somewhat to

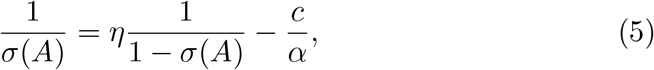

And

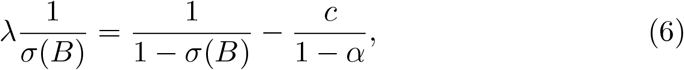

respectively. These are quadratic equations that have the following unique closed form solution (in the relevant range between 0 and 1).

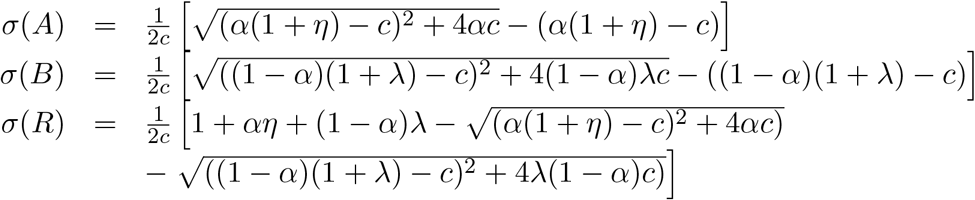

If, moreover, we consider the limit case of cognition costs going to zero, *c* → 0, the system of equations further simplifies to

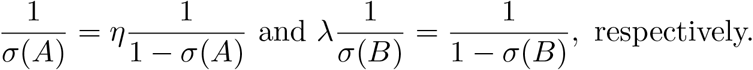

The unique solution is then 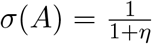 and 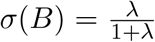 this is the only candidate for an equilibrium of this form. As *σ*(*R*) = 1 − *σ*(*A*) − *σ*(*B*) we obtain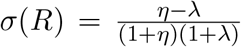. Note, finally, that *σ*(*A*), *σ*(*B*), *σ*(*R*) ≥ 0, and, thus *σ* is indeed the unique (mixed strategy) equilibrium.

## 4 Equilibrium Implications

Note first that the equilibrium frequency *σ*(*A*) of always going to the constant food source *A* does not depend on *λ*, the low state of food-availability in the stochastic source: *σ*(*A*) does not change when *λ* changes. This is not only true in the case of *c* → 0 and *n* → *∞*, but generally true. This can be seen by the fact that Equation 3 is an equation just in *σ*(*A*) (not also in *σ*(*B*) or *σ*(*R*)) and the parameter *λ* does not appear in this equation. Similarly, the equilibrium frequency *σ*(*A*) of always going to the stochastic food source *B* does not depend on *η*, the high state of food-availability in the stochastic source. Only the equilibrium frequency, *σ*(*R*) of being responsive changes with both levels of food-availability of the stochastic source. For a fixed high level of food-availability *η* at the stochastic source, increasing *λ* leaves *σ*(*A*) constant, increases *σ*(*B*), and decreases *σ*(*R*), the frequency of responsive individuals. Analogously, for a fixed low level of food-availability *λ* at the stochastic source, increasing *η* decreases *σ*(*A*), leaves *σ*(*B*) constant, and increases *σ*(*R*). All these findings are illustrated in Figure 2. If both *η* increases and *λ* decreases (i.e., the stochastic food source becomes more extreme) then *σ*(*A*) and *σ*(*B*) decrease and, therefore, the equilibrium frequency of responsive individuals, *σ*(*R*) increases.

**Figure 2:**
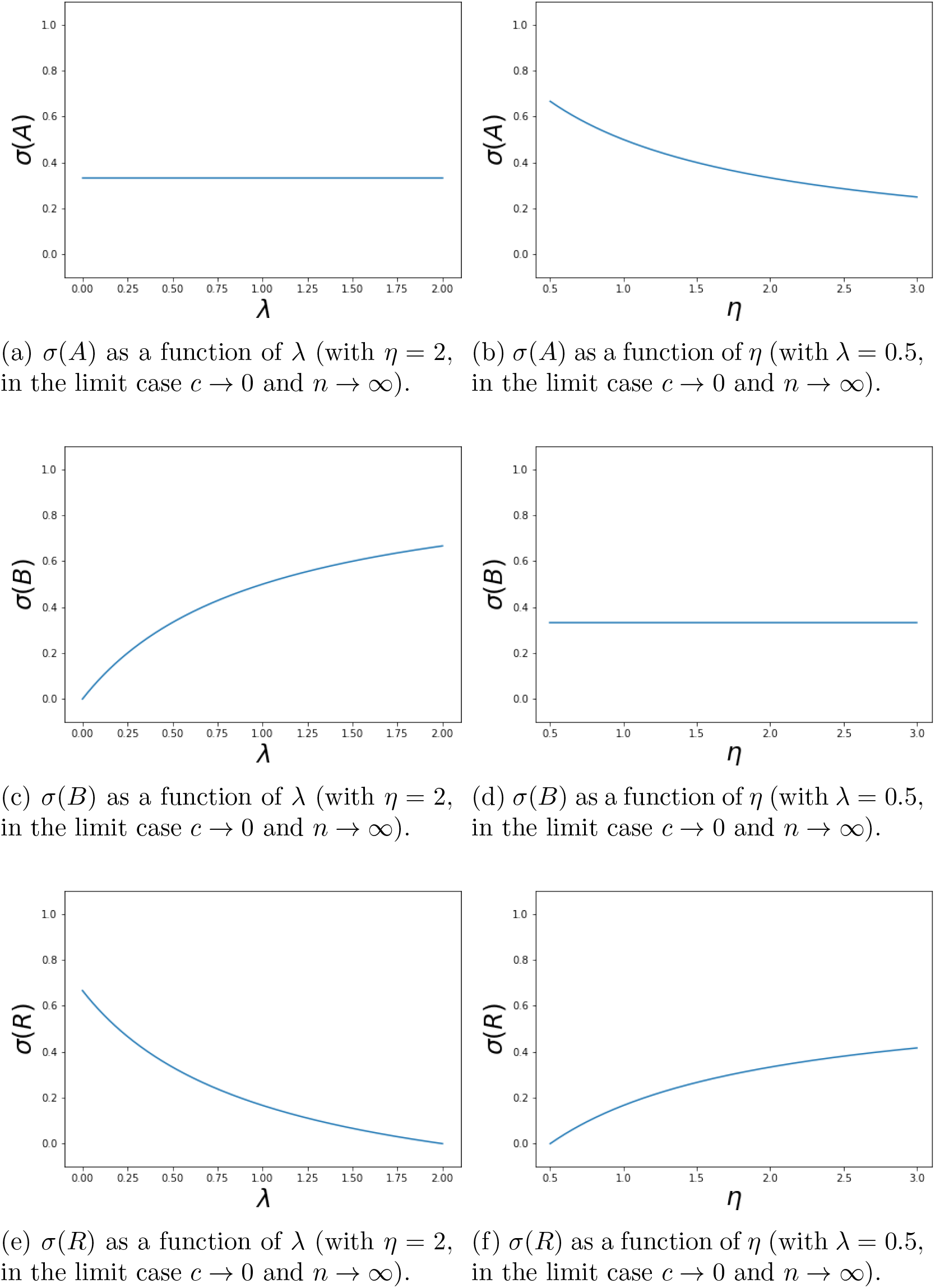

The second thing to note is that in the limit that the cost of cognition becomes negligible (relative to the other parameters), that is when *c* → 0 (for any *n*), the equilibrium frequencies do not depend on the stochastic nature of the environment, i.e., do not depend on *α*. This can be seen by setting *c* = 0 in equations 3 and 4, with the result that *α* drops out of these equations. If the cost of cognition, *c*, is non-negligible, then the equilibrium frequencies depends on this cost and on *α*, the stochastic nature of the environment, as illustrated in Figure 3.

**Figure 3:**
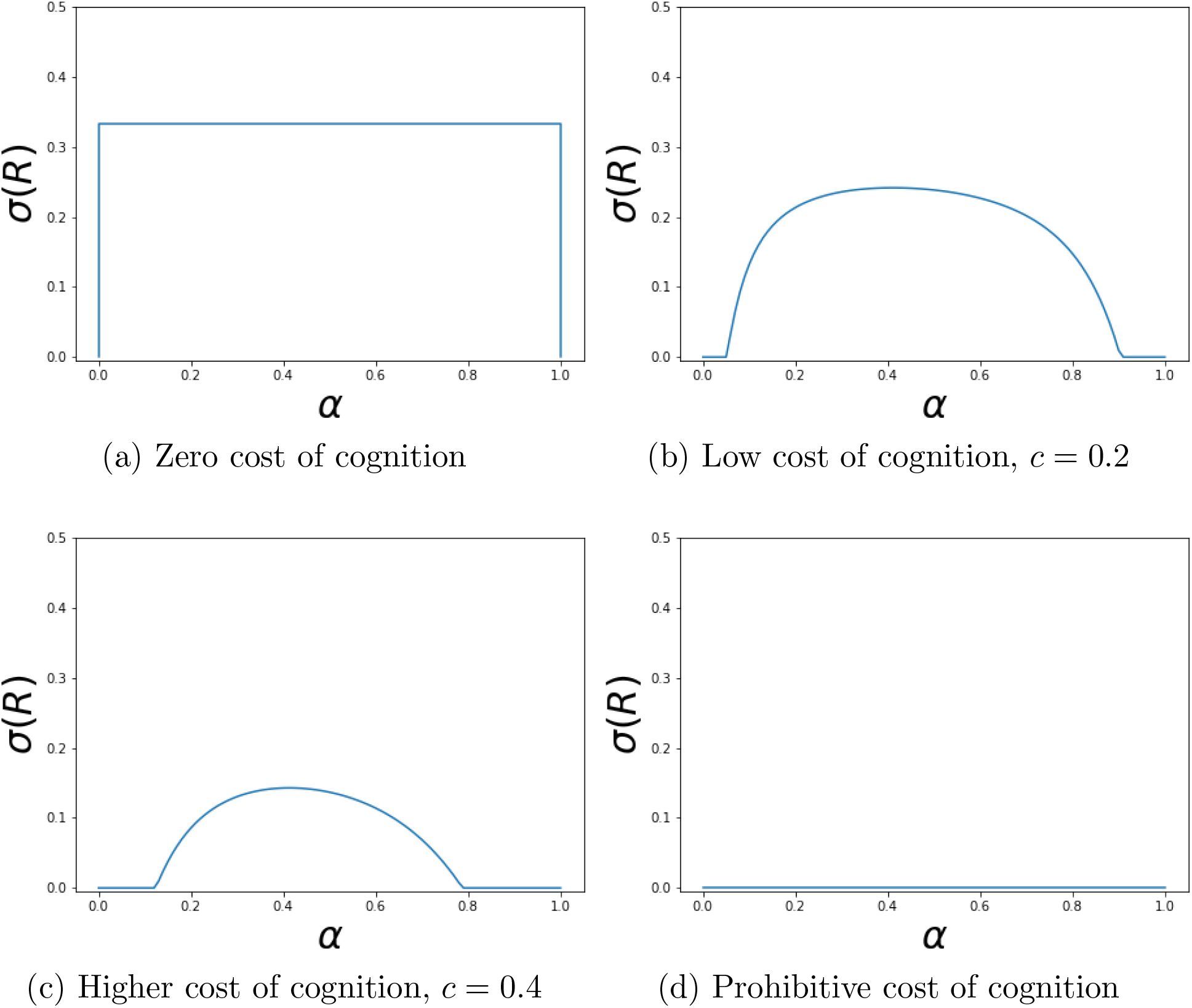
Probability or frequency of being responsive, *σ*(*R*) as a function of the probability, *α* of the high food availability at the stochastic food source, for different levels of the cost of cognition, *c* (*λ* = 0.5, *η* = 2).

In Appendix C we show that the unique symmetric Nash equilibrium in this model is also an evolutionarily stable strategy, in the sense of Palm [1984], see also Broom et al. [1997], who have extended the definition of an ESS of Maynard Smith and Price [1973] to symmetric *n*-player games. We know generally, see e.g., Nachbar [1990], that Nash equilibria are the only candidates for asymptotically stable rest points under most deterministic behavioral adjustment (or evolutionary) dynamics, with the replicator dynamics of Taylor and Jonker [1978] the first and most prominent example. In Appendix D we show that the games we here study are stable games in the sense of Hofbauer and Sandholm [2009], with the implication that the unique symmetric equilibrium of our model is asymptotically stable under many different dynamics that includes the replicator dynamics. In Figure 4 we illustrate this finding by sketching the phase diagram of the replicator dynamics for different parameter configurations of our model.

**Figure 4:**
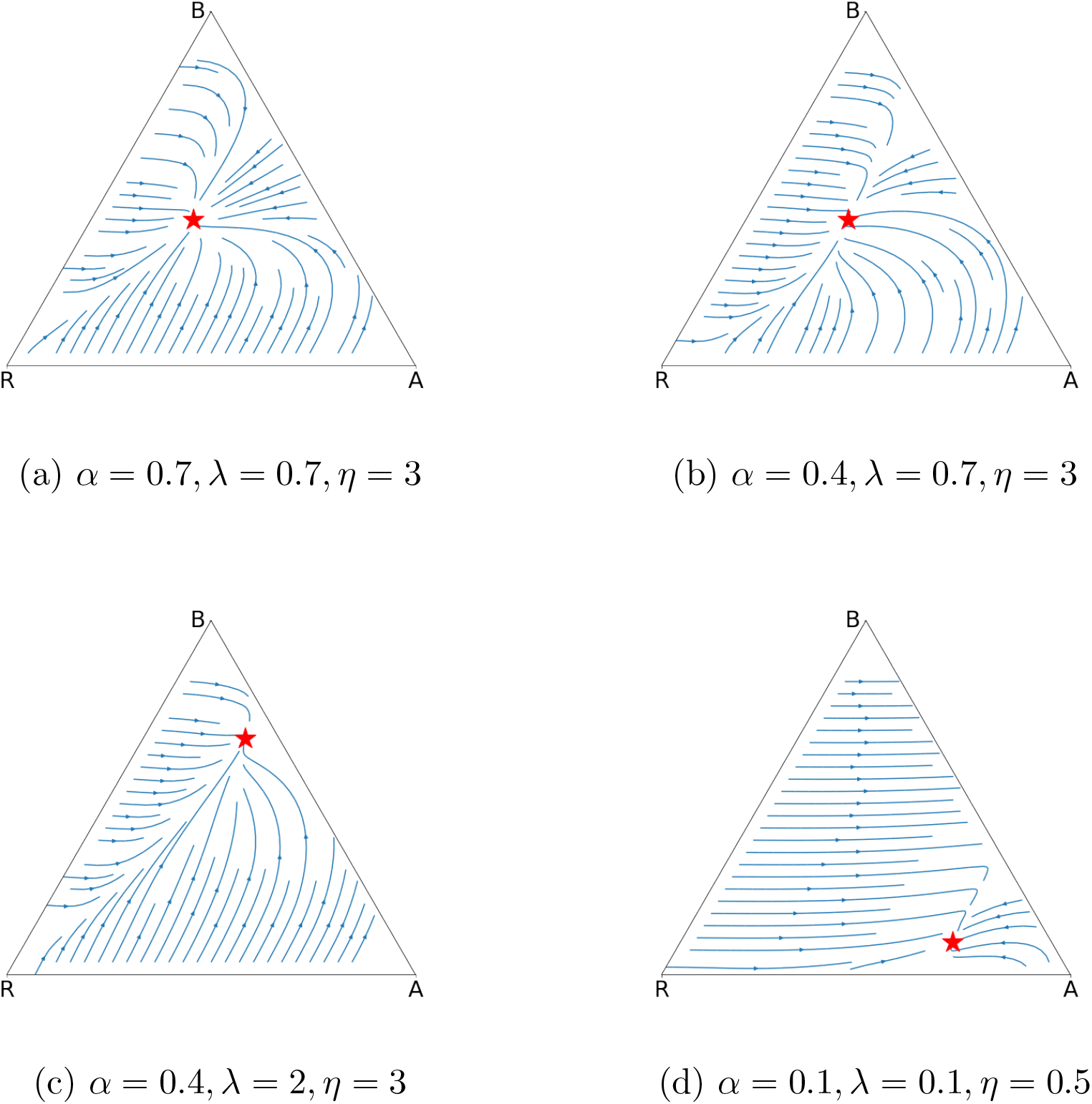
Phase diagram of the replicator dynamics for *c* = 0, under a variety of different parameter specifications. The only difference between figures *a* and *b* is that the stochasticity parameter *α* changes. This has no effect on the equilibrium itself but does affect somewhat the out of equilibrium dynamics. It can also be seen that not only is the unique equilibrium asymptotically stable, but also in fact a global attractor under the replicator dynamics: All solution paths eventually converge to the equilibrium.

## 5 Consistency and Correlation

Suppose that the *n* individuals play the same game given in our basic model over and over again for many periods of time. Suppose that, at every point in time, they play the unique symmetric equilibrium given earlier. An outside observer would note that when the amount of food available at *B* is high (*X* = *η*) more individuals are to be found at source *B* than when it is low (*X* = *λ*). They would also observe that the food share each individual receives is the same regardless of which source the individuals go to. The outside observer would conclude that some individuals must be responsive to the stochastic food availability at source *B*. However, there are, so far, no incentives for individuals to choose the same strategy across time and context.

Recall the two interpretations we can give a mixed strategy equilibrium, as we identified in this model. Suppose that we interpret it as the various individuals involved actually randomizing between the three pure strategies. In that case, the outside observer would note that the identity of responsive individuals varies over time. This, however, is inconsistent with empirical findings (of consistency and correlation as highlighted in the introduction). Alternatively we can interpret the mixed strategy equilibrium as there being a large population of individuals, with certain fractions of these individuals playing the various pure strategies. In this latter case, it is more plausible that if one individual is responsive at one point in time it is also responsive at another point in time. Note, however, that all individuals only have a very weak incentive to stay with their pure strategy choice, as (in any mixed equilibrium) all pure strategies provide the same expected benefit (food share in our model). In principle, the various individuals could switch pure strategies around, as long as the aggregate frequencies remain the same. A slight change to the basic model, however, provides essentially the same prediction as the basic model, but now with every individual having a strict preference to play a pure strategy with frequencies very close to the mixed strategy equilibrium of the basic model.

This modification is based on the idea of *purification* of Harsanyi [1973], which is very similar to the idea of *threshold decisions* as provided in McNamara and Houston [2005]. The idea is that individuals differ a little bit in terms of their personal preferences and actually make a pure strategy choice that is, however, dependent on their own personal preferences that only they themselves know. As a consequence, while the equilibrium looks mixed to other individuals, each individual actually plays a pure strategy.

We adapt the model by replacing the payoff function *u* of the original model with a slightly perturbed payoff function *v*_*θ*_ that is essentially equal to *u* plus a small idiosyncratic (individual-specific) preference or perturbation term:

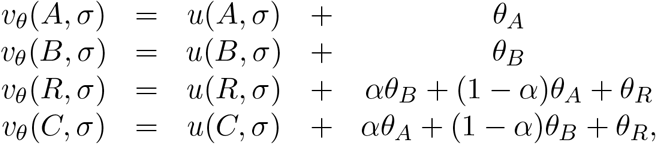

where the vector *θ* = (*θ*_*A*_, *θ*_*B*_, *θ*_*R*_) is i.i.d. drawn from some arbitrary full support continuous joint distribution *F* (with density *f*) over Θ = [−*ϵ, ϵ*]^3^, for a small *ϵ >* 0. It is assumed that an individual’s realized *θ* is that individual’s private information, which means that only this individual knows it; it is unknown to other individuals. We have deliberately chosen the same preference perturbation *θ*_*R*_ for pure strategies *C* and *R*, as it seems more natural to have an idiosyncratic perturbation of the cost of being responsive rather than for how one is responsive. However, it does not matter what we assume for pure strategy *C* as long as the payoff perturbation is small, as pure strategy *C* provides a strictly lower payoff than the other three strategies in the equilibrium given earlier, and small payoff perturbations cannot change that.

In the perturbed game, a strategy is a function *ρ* : Θ → Δ(*S*), where *ρ* attaches to each possible preference type *θ* a probability distribution over *S*. Every such strategy *ρ* induces a strategy *σ*^*ρ*^ ∈ Δ(*S*) by *σ*^*ρ*^(*s*) = ∫ _Θ_ *ρ*(*θ*)(*s*)*f* (*θ*)*dθ* for all *s* ∈ *S*, where *ρ*(*θ*)(*s*) is the probability that *ρ*(*θ*) attaches to *s*. An equilibrium strategy *ρ* is then such that *ρ*(*θ*)(*A*) = 1 if

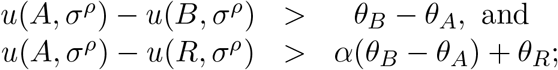

*ρ*(*θ*)(*B*) = 1 if

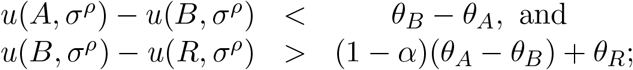

and *ρ*(*θ*)(*R*) = 1 if

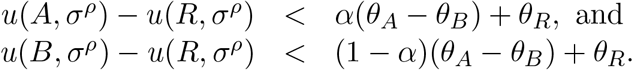

Harsanyi [1973] has shown that almost any equilibrium of a complete information game, such as our basic game, is such that any nearby incomplete information game, with payoff perturbations given by the joint distribution *F*, has a nearby equilibrium and that this nearby equilibrium is essentially in pure strategies. In such a nearby equilibrium there is a parameter region for *θ* ∈ Θ for which an individual strictly prefers to play *A*, another region for which an individual strictly prefers to play *B*, and a final region in which an individual strictly prefers to play *R*. The set of *θ*’s for which an individual is indifferent between two or three of the three strategies has measure zero (it is statistically impossible). Finally, such purified equilibria can also shown to be dynamically stable under a suitably defined behavioral adjustment dynamics as in Ely and Sandholm [2005], see Sandholm [2007]. In other words, the result of Harsanyi [1973] implies that each individual uses a pure strategy, which they strictly prefer given their own private preference, but the frequency of each strategy at the population level is essentially the same as without the preference perturbation.

Suppose now that the *n* individuals play the resulting equilibrium of the same slightly perturbed game repeatedly over many time periods. It is then a question of whether the perturbation parameters *θ* remain the same for each individual over time or not. If they do, it will be the same individuals who always go to food source *A*, the same individuals who always go to food source *B*, and the same who are responsive.

As an example of why the perturbed model may be appropriate for our purposes, consider birds who every day have to decide to go to food source *A* or *B* from their nesting place. Then the location of their nesting place gives rise to their *θ*. An approach could be that *θ*_*A*_ and *θ*_*B*_ are proportional to the distance that the bird’s nest is from the two food sources, respectively, while *θ*_*R*_ could be more of a personal characteristic of the bird, measuring how much/less cognitively able this bird is relative to other birds.

This model is then also flexible enough to generate a strong consistency over time and a weaker, but some, consistency across contexts, depending on how these consistencies are interpreted. Consider the bird example again. One could imagine that *θ*_*R*_ is an individual bird’s specific parameter that does not change over time nor across contexts. On the other hand, the parameters *θ*_*A*_ and *θ*_*B*_ might be constant for one season, but could be different in another season, when the bird’s nest location (or the location of the food sources) changes.

Equilibrium purification could even be obtained by introducing a payoff-irrelevant personal and privately know characteristic, such as an individual’s prior experiences in life, with individuals playing different pure strategies depending on their personal prior experiences. This means that, as pointed out e.g., in Wolf et al. [2007], Wolf and Weissing [2010] and Dingemanse and Wolf [2010], the purification threshold could also be based on an individual’s life history.

## 6 Imperfect private signals of food availability

In our basic model, individuals can learn the state of food availability at the food sources perfectly. In this section, we study how the results change if this learning is imperfect. To do so, we suppose that each individual receives a noisy signal about the level of food availability at food source B. Individuals *i* receive conditionally independent (and identically distributed) signals *s*_*i*_ ∈ *{l, h}* such that *P* (*s*_*i*_ = *h*|*X* = *η*) = *P* (*s*_*i*_ = *l*|*X* = *λ*) = 1 − *ϵ*, With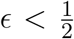. In words, in the high state *η* the high signal *h* is more likely than the low signal *l*, and in the low state *λ* the low signal *l* is more likely than the high signal *h*. The signal is, thus, informative about the true state. All the arguments made for the perfect signals of the food availability model (the basic model) go through (see Appendix B). Ultimately, we obtain that, provided the level of noise *ϵ* is not too large, the game has a unique symmetric Nash equilibrium, denoted *σ*_*ϵ*_, that is also an ESS (and dynamically stable under a large class of evolutionary adjustment models). In the limiting case when the cost of cognition *c* → 0 the equilibrium is given by

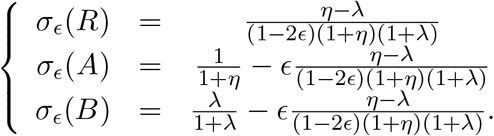

There are two interesting implications of this model, both seen also in Figure 5. First, for all parameter values *λ, η*, the frequency of responsive individuals increases with the level of noise *ϵ* in the information. The noisier the signal the more individuals are responsive to the possible erroneous information about the stochastic food source. The intuition behind this is, that because of the possible error, more individuals have to respond, because some will get the wrong signal and, thus, respond incorrectly. This would lead to not enough people responding appropriately. Second, the impact that an increase of the level of noise *ϵ* has on the frequency of responsive individuals is higher the higher the frequency of responsive individuals is without noise. This can be seen in Figure 5, and also formally, if we write

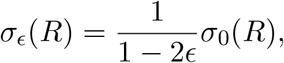

and, taking the derivative with respect to *ϵ* obtain

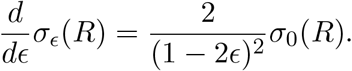

Interestingly, these results remain valid even when considering a strictly positive cognition cost.

**Figure 5:**
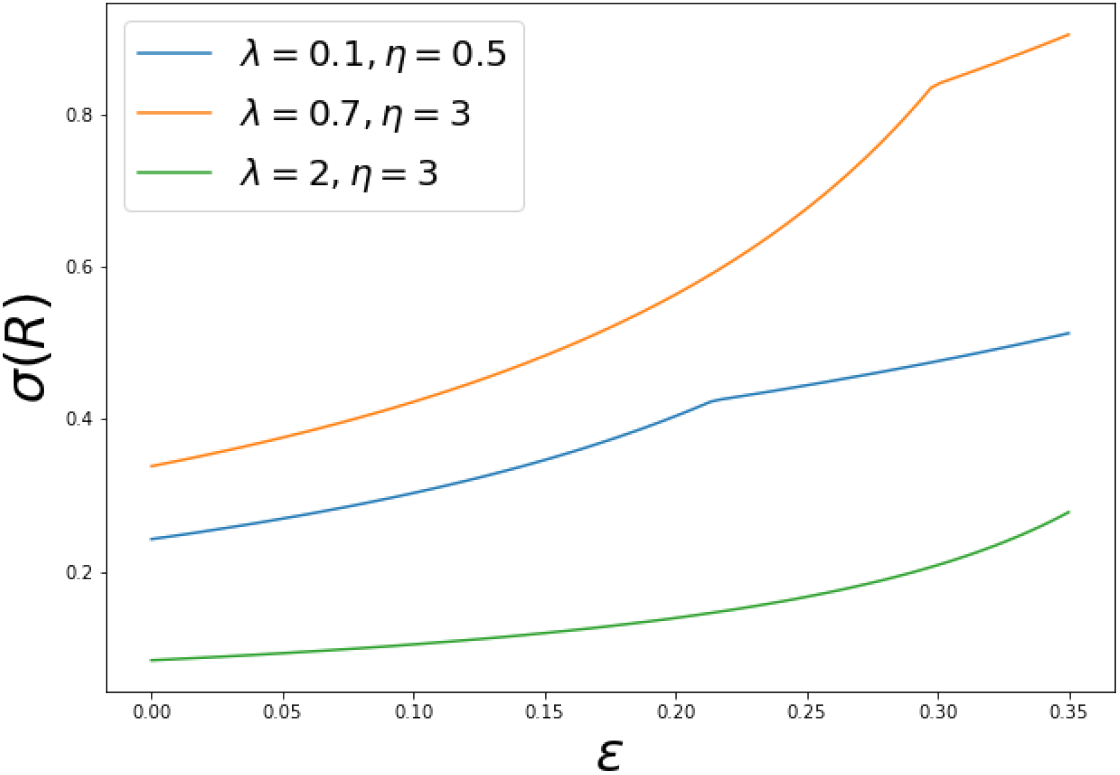
The proportion of responsive individuals increases with the level of noise in the information. The kink in the graph is at the level of noise *ϵ* at which one of the two equilibrium frequencies *σ*_*ϵ*_(*A*) or *σ*_*ϵ*_(*B*) become zero.

## 7 General distributions of food source availability

Recall that in the basic model food source B is assumed to be Bernoulli distributed (i.e., with only two possible levels of available food at that source). In this section we consider an arbitrary distribution for the food availability at food source B.

Let *X*, the available quantity of food at food source B, be distributed according to some distribution with cdf *F* with everywhere positive density *f* on the interval *χ* = [*x*_*L*_, *x*_*H*_] with 0 ≤ *x*_*L*_ *< x*_*H*_ ≤ *∞*. To make the analysis tractable we simplify the model in two ways. First, we set the cost of being responsive, *c*, to zero.^7^ Second, we assume that all individuals learn the value of *X*, and allow individuals to only use monotone strategies: An individual’s strategy can be described by a cutoff value *y* ∈ *χ* such that the individual goes to food source A if and only if *x < y*. Otherwise the individual goes to food source B. This implies that the strategy space is identical to *χ* and the set of mixed strategies is the set Δ(*χ*) of all probability distributions over *χ*. A fully mixed symmetric Nash equilibrium strategy, which can be described by a cdf *G* on *χ* must satisfy that any individual is indifferent between using any any cut-off *y* ∈ *χ*.

We then get the following result. In the model of this section, for any *n*, there is a unique completely mixed symmetric equilibrium. In the limit as *n* tends to infinity, the equilibrium probability that an individual uses cut-off responsiveness *y* is given by the cdf 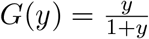, with 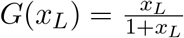 the probability of an individual always going to food source B, 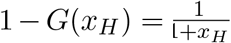 the probability of an individual always going to food source A, and 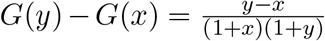 the probability that an individual adopts a degree of responsiveness in the interval [*x, y*].^8^

This finding is consistent with those in the basic model. For example, the strategy called *B* in the previous model is similar to choosing the cut-off *x*_*L*_ (since *x*_*L*_ is the minimum possible value of the stochastic source). The equilibrium frequency of this strategy in the basic model is given by 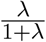, which is equivalent to 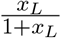, as *x*_*L*_ is the smallest possible value of the stochastic source. This generalization to the basic model also delivers a new insight. If there is a wide range of possible levels of food availability at the stochastic food source, then in equilibrium there is a continuum of degrees of responsiveness to environmental stimuli. See e.g., Lionetti et al. [2018] for empirical support for this finding.

## 8 Discussion

We built a stylized game-theoretic model of foraging behavior in a stochastic environment. For every parameter specification within certain bounds, this model has a unique symmetric Nash equilibrium, that is also the unique ESS and asymptotically stable under a variety of evolutionary dynamics. This equilibrium has the three key features identified in the literature of *coexistence* of differing degrees of environmental responsiveness, *consistency* of individual responsiveness over time, and *correlation* of individual responsiveness across contexts.

By explicitly studying the phenomenon of heterogeneous responsiveness to environmental stimuli in a foraging setting, we are able to identify the push towards the ideal free distribution of Fretwell and Lucas [1969], satisfied in the equilibrium of our game, as a possible driving force of this heterogeneity. See also Křivan et al. [2008] for a game-theoretic treatment (without environmental stochasticity). We derive explicit analytical expressions for the equilibrium frequencies of responsive and non-responsive behavior, at least when the cost of cognition (needed to respond to environmental stimuli) is negligible or at least relatively small. This allows us to study how the equilibrium frequencies change when some of the model parameters change. For instance, we find that, at least when cognition costs are negligible, the exact stochastic nature of the environment does not affect the equilibrium. This finding suggests that changes in the stochastic environmental process would at least not be so disruptive as to push behavior out of equilibrium. Put differently, equilibrium strategies are already complex enough to allow for automatic adaptation to such changes in the stochastic environmental process. In the remainder of this section we discuss some of the limitations of our approach.

### Cost of cognition

We have mostly explored the case of zero, and by a continuity argument, also of small cost of cognition. For larger cost of cognition, generally, equilibrium behavior will depend on the stochastic nature of the environment, see, for instance, Figure 3, and the equilibrium will not satisfy the ideal free distribution. For most species, however, it is not unreasonable to assume a relatively small cost of cognition, see e.g., [Hendry, 2016, Murren et al., 2015, Auld et al., 2010, Van Buskirk and Steiner, 2009]. Relatedly, the cost of plasticity, as defined, for instance, in DeWitt et al. [1998], see also [Relyea, 2002] is typically small; see, for instance Hendry [2016], the review by Murren et al. [2015], and the meta-analysis by Van Buskirk and Steiner [2009].

### Noisy information

Another insight that we can derive from an extension of our model is that the higher the noise in the environmental stimuli the more responsive individuals become in equilibrium. This is under the assumption of individuals receiving private and stochastically independent noisy information about the state of the environment. We have not explored the case of correlated information, such as all individuals receiving the same public information. In such a setting, the ideal free distribution would at best hold in expectation, and there would be a positive variance of food share availability at the random source.

### Social Information

Another, empirically relevant, informational setting, that we here abstracted away from, is one where not all individuals receive the same quality of information (perhaps not all are equally close to the source of information). One would then expect individuals to infer additional information about the state of the environment from other individuals’ behavior. If, for instance, there are many birds flying out to a specific point at sea, another bird might follow them based on the idea that there is information in that behavior. This will certainly be the case for socializing birds, which display behavior of forming flocks and swarms. Such behavior would add another layer of complexity to the game. For a literature review on social information use in a foraging context see Kohles et al. [2022]. According to the “information-sharing theory”, as in Clark and Mangel [1984], it has been observed for different species that animals can both search for food on their own or join others who have found food, see e.g., Giraldeau and Beauchamp [1999]. Models, in which individuals choose between the two strategies “discoverers” or “copiers” are referred to as “producer-scrounger games” [Giraldeau and Beauchamp, 1999, Bhattacharya and Vicsek, 2014]. The interest in using social information is notably to avoid paying a cost to acquire private information [Webster and Laland, 2008]. The choice between producer and scrounger is, to some extent, persistent in time [Morand-Ferron et al., 2011]. The connection between *responsiveness* and the producer-scrounger game could be made through the concepts of *personality*. Indeed, the main trait of personality is the axis boldness-shyness, and it has been observed that shyness increases the probability to scrounge [Kurvers et al., 2010].

### The number of food sources

Our model only has two food sources. This keeps the analysis mathematically tractable, but at the cost of a possible oversimplification. Given our results, however, one would conjecture that in any (evolutionary stable) equilibrium of a game with multiple food sources, the ideal free distribution holds, at least when costs of cognition are negligible: all food sources would have equal food shares, and this would be true for all states. This would imply that equilibrium behavior would not depend on the exact stochastic nature of the environment. This would also imply that any (evolutionary stable) equilibria would again satisfy coexistence, and for slightly perturbed models, consistency and correlation. One would, however, not necessarily expect a unique equilibrium and it would be harder to characterize these explicitly.

### Generalizing

We focused here on foraging choice as a concrete setting in which one would expect the coexistence, consistence and correlation of different responsiveness to external stimuli. However, our results could possibly be generalized to different context where there is a resource to share among individuals, with a resource distributing at different point, and some of them stochastic. Such contexts are social interactions, mating behaviour, division of labour [Dall et al., 2012], space-use [Spiegel et al., 2017, Ceia and Ramos, 2015], or niche specialisation [Dall et al., 2012, Montiglio et al., 2013, Bolnick et al., 2003]. Those last studies show similar concepts (state dependence, frequency dependence, social awareness, environmental heterogeneity) applied to niche specialisation. In particular, increasing evidence show a link between specialisation and personality. It is hypothesized that personality implies specialization [Toscano et al., 2016, Harris et al., 2020] or the other way around [Bergmüller and Taborsky, 2010, Montiglio et al., 2013]. Similar to our model, Leimar et al. [2022] used frequency-dependent competition for resources to explain specialization. For a review of individual foraging specialization see Sheppard et al. [2021]. Finally and more speculatively, our results might even be translatable to the issue of stem cell differentiation, which might arise from competition over resources, see Jörg et al. [2019].

## A General distributions of food source availability (proof)

Let *u*(*A, G, x*) and *u*(*B, G, x*) denote the expected payoff to an individual who goes to food source A and B, respectively, when all others use mixed strategy *G* and when the realized state of food availability at food source B is given by *x*. As *G*(*x*) = *P* (*y* ≤ *x*) is the probability that a random other individual goes to food source B, using again [Chao and Strawderman, 1972, Equation 3.4], we have

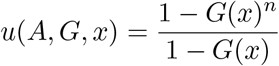

And

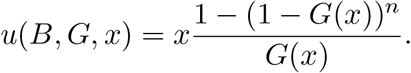

The expected payoff for an individual using cut-off *y* ∈ *χ*, when all others use mixed strategy *G*, is then given by

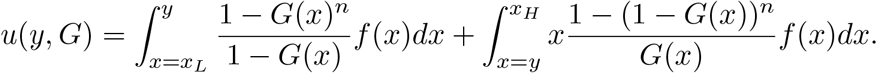

For a completely mixed strategy *G* to be a symmetric Nash equilibrium we need that *u*(*y, G*) is constant for all *y* ∈ *χ*. If *u*(*y, G*) is a constant function in *y* then its derivative with respect to *y* must be constant as well (and equal to zero). This implies that

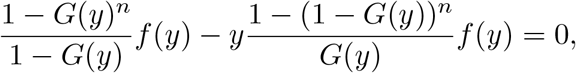

for all *y* ∈ *χ*. Note that *f* (*y*) cancels out and the resulting equilibrium distribution *G* does again not depend on the stochasticity of the food source, and we obtain

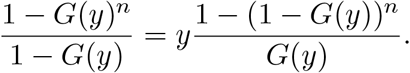

Note that the left hand side of this equation is increasing in *G*(*y*) and the right hand side is decreasing in *G*(*y*). Note also that the left hand side is strictly less than the right hand side when both are evaluated at *G*(*y*) = 0 and the inequality is reversed when both are evaluated at *G*(*y*) = 1. Thus, for any *y* ∈ *χ*, this equation has a unique solution *G*(*y*). Note that this solution *G*(*y*) must necessarily be higher for higher *y*. Note also that, necessarily, *G*(0) = 0 and lim_*y*→*∞*_ *G*(*y*) = 1. *G* is, thus, a cumulative distribution function.

For finite *n* this solution is not easily expressed analytically. In the limit as *n* → *∞* we obtain

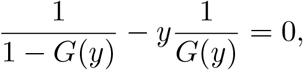

for all *y* ∈ *χ*, and, thus, 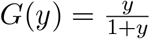 in this limiting case.

## B Imperfect signals

In this case, (random) food shares are given by

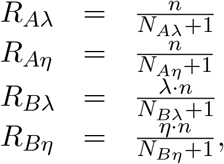

where, given strategy *σ* (used by all opponents), *N*_*Aλ*_ follows a binomial distribution with *n* − 1 trials and success probability *σ*(*A*) + (1 − *ϵ*)*σ*(*R*) + *ϵσ*(*C*). Similarly, *N*_*Aη*_, *N*_*Bλ*_, *N*_*Bη*_ are also binomially distributed with *n* − 1 trials and success probabilities *σ*(*A*) + (1 − *ϵ*)*σ*(*C*) + *ϵσ*(*R*), *σ*(*B*) + (1 − *ϵ*)*σ*(*R*) + *ϵσ*(*C*), and *σ*(*B*) + (1 − *ϵ*)*σ*(*C*) + *ϵσ*(*R*), respectively.

We, thus, obtain

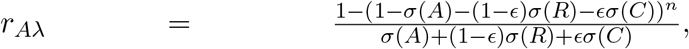

and analogously

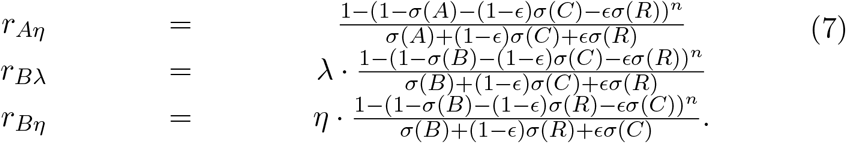

We need to show that there is a unique equilibrium with support *{A, B, R}*. To be indifferent between *A* and *R* requires that

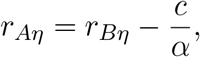

to be indifferent between *B* and *R* requires

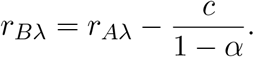

Plugging in the above expressions, and noting that *σ*(*C*) = 0, these two equations lead to

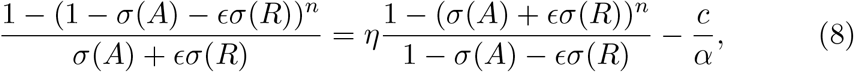

And

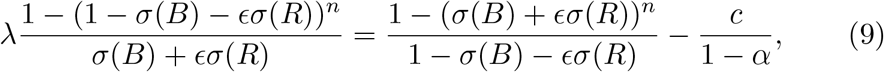

respectively. We are interested in the limit as *n* → *∞* and *c* → 0 and obtain

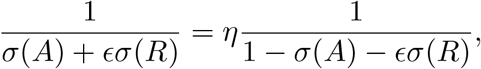

and

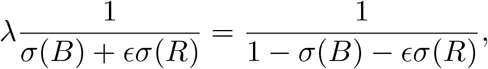

respectively. Solving these two equations yields unique solutions 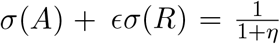 and 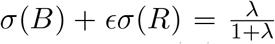 equilibrium of this form. As *σ*(*R*) = 1 − *σ*(*A*) − *σ*(*B*) we obtain 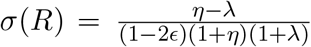and then 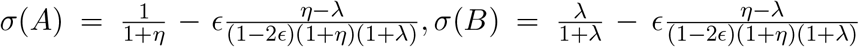 . Note, finally, that *σ*(*A*), *σ*(*B*), *σ*(*R*) ≥ 0, and, thus *σ* is indeed the unique (mixed strategy) equilibrium.

When *ϵ* gets large enough, one of the nonreactive strategies *A* or *B* can vanish. Then, the equilibrium changes. For example, if *B* vanishes. Then, the only equation is 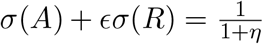 with *σ*(*A*) + *σ*(*R*) = 1, which give the equilibrium 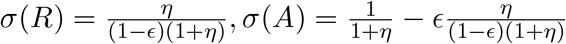

## C Evolutionary Stable Strategy

We here show that the unique symmetric equilibrium of the basic game is evolutionarily stable as defined, for symmetric *n* player games, by Palm [1984], see also Broom et al. [1997]. By [Palm, 1984, Proposition 4] a sufficient condition for a (mixed) strategy *σ* of a symmetric *n*-player normal form game is an ESS is that

1. *u*(*σ, σ*) ≥ *u*(*σ*^*′*^, *σ*) for all *σ*^*′*^ ∈ Δ(*S*) and
2. if *u*(*σ*^*′*^, *σ*) = *u*(*σ, σ*) for some *σ*^*′*^≠*σ* then *u*(*σ*^*′*^, *σ*^*′*^) *< u*(*σ, σ*^*′*^).

The first condition states that only symmetric Nash equilibria can be an ESS. Thus, the strategy we identified as the unique symmetric Nash equilibrium of the game is the only candidate for an ESS. To show that it is indeed an ESS we need to show that it also satisfies the second condition.

For strictly positive costs pure strategy *C* yields a strictly lower payoff than the equilibrium payoff. However, all strategies *σ*^*′*^ ∈ Δ(*S*) with *σ*^*′*^(*C*) = 0 satisfy *u*(*σ*^*′*^, *σ*) = *u*(*σ, σ*). We need to show that for all such *σ*^*′*^≠*σ* we have *u*(*σ, σ*^*′*^) − *u*(*σ*^*′*^, *σ*^*′*^) *>* 0. As *σ*^*′*^(*C*) = *σ*(*C*) = 0. We have that,

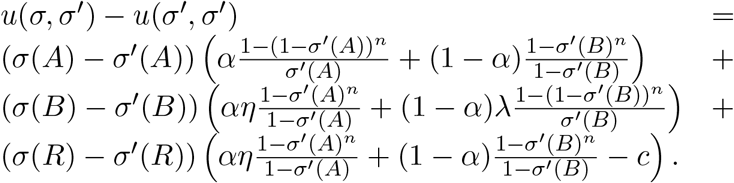

As *σ*(*R*) = 1 − *σ*(*A*) − *σ*(*B*) (and the same for *σ*^*′*^), this is equivalent to

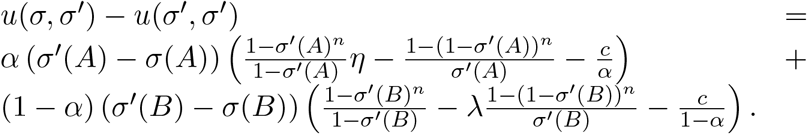

Note that last term in brackets in the first line is the equilibrium equation 5 (here for *σ*^*′*^(*A*)) and similarly the last term in brackets in the second line is the equilibrium equation 6 (here for *σ*^*′*^(*B*)). These terms are, therefore zero, for *σ*^*′*^ = *σ* and they are, for all *n* ≥ 2, increasing functions in *σ*^*′*^(*A*) and *σ*^*′*^(*B*), respectively. To see this for the first of these two expressions,

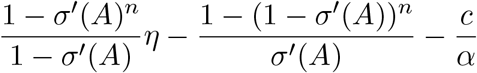

note that the first term in this expression is

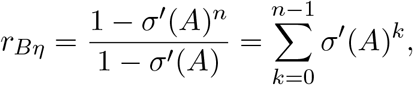

and, thus, clearly increasing in *σ*^*′*^(*A*). Analogous arguments can be made for the second term in this first expression, as well as for both terms in the second of the two expressions. This means that the first term in brackets is negative exactly when *σ*^*′*^(*A*) − *σ*(*A*) is negative and is positive exactly when the latter is positive. The same argument is true for the second term in brackets. Therefore, for any *σ*^*′*^≠*σ* with *σ*^*′*^(*C*) = 0 we, indeed, have *u*(*σ, σ*^*′*^) − *u*(*σ*^*′*^, *σ*^*′*^) *>* 0. Therefore, the equilibrium *σ* is an ESS.

## D Dynamic stability

In this section we show that the class of games we here study are all stable games, in the sense of Hofbauer and Sandholm [2009]. Results in Hofbauer and Sandholm [2009] then imply that the unique symmetric Nash equilibrium that we have identified above is asymptotically stable under a large class of evolutionary (or behavioral adjustment) dynamics, that includes, for instance, also the well-known replicator dynamics of Taylor and Jonker [1978].

In order to prove that our games are stable games, we need to introduce some of the more general notation introduced by Hofbauer and Sandholm [2009]. As we have just a single populations of players, in their notation we have *p* = 1. In their notation *X* denotes the space of players’ mixed strategy space, which in our setting is, therefore, equal to Δ(*S*). The payoff function in their setting is denoted by *F* : *X* → **R**^*k*^, where *k* is the number of pure strategies, i.e., *k* = 4 in our setting.^9^ Translated to our notation, *F* is, therefore, given by

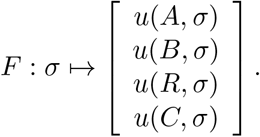

A population game *F* : *X* → **R**^*k*^ is a stable game, as defined by Hofbauer and Sandholm [2009], if,

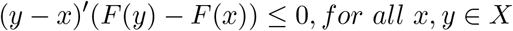

We now prove that our games are stable games in this sense. We utilize [Hofbauer and Sandholm, 2009, Example 2.9] to show that our games are *negative dominant diagonal games*, which by [Hofbauer and Sandholm, 2009, Proposition 2.7] are stable games.

To do so we need to check two conditions. First, a condition of universal *negative frequency dependence*,

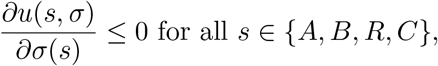

and second,

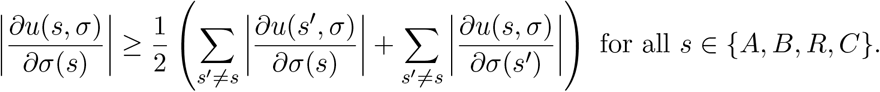

We here only provide the proof that these two conditions are satisfied for *s* = *A*, the proof for other strategies in *S* = *{A, B, R, C}* proceeds analogously.

Recall that,

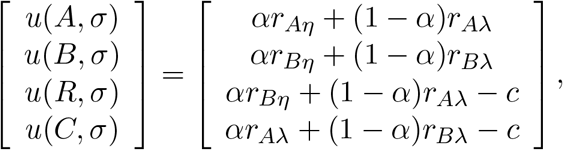

where, by Equation 2,

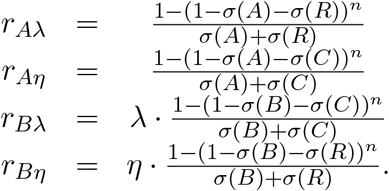

We, therefore, have that

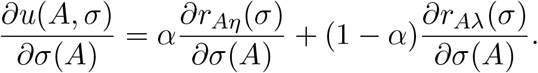

Note that

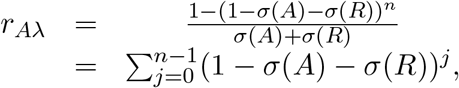

by the well-known formula for the geometric series. Thus,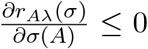. By the same argument 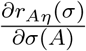 is also everywhere non-positive, and, therefore,

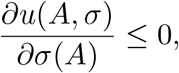

as desired. Analogous arguments yield that the desired first condition is satisfied:

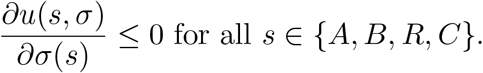

Let us now consider the second condition, again for *s* = *A*, which can be written as

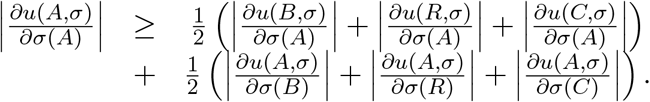

We prove this statement in two steps. Note first that

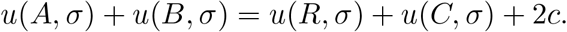

Therefore,

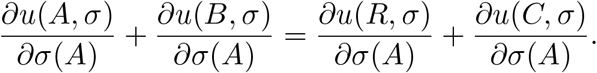

Since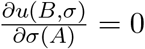, we then have

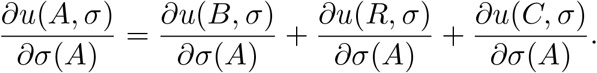

As all these derivatives are either negative or equal to zero, we have

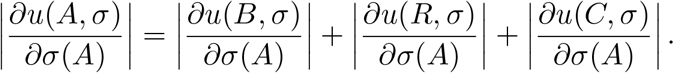

For the second step, note that 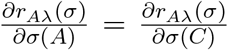, as well as 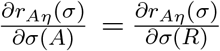 . Moreover, because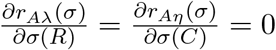, we have,

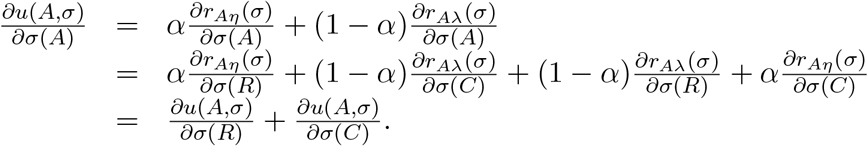

As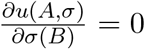, and again, as all derivatives are either negative or equal to zero we have

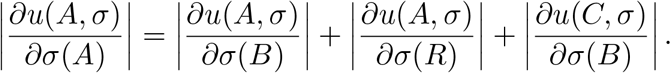

Finally, by adding the result of the two parts, the second condition follows for *s* = *A*. For strategies *B, R, C* the same result can be proven in an analogous manner.

## Footnotes

1 Stamps [2016] refers to “[t]he extent to which the phenotype of an agent varies as an immediate response to variation in external stimuli” as “contextual plasticity.”

2 The “suite of correlated behaviors […] reflecting the individual consistency across […] situations” has been referred to as a “behavioral syndrome” by Sih et al. [2004].

3 Pluess [2015], also Mitchell and Houslay [2021], argue in favor of an integration of the two theories.

4 Wolf et al. [2008], Ehlman et al. [2022] explain consistency and correlation with a “positive-feedback mechanism:” responsiveness is less costly for individuals that have been responsive before. Wolf and McNamara [2012] explain consistency and correlation by small variations of individuals’ metabolism (which is a form of state).

5 In principle, they could also become informed and then ignore their information, by going to *A* or *B* regardless of the information they have received. But we assume that the choice of becoming informed bears an arbitrarily small cost *c >* 0. The assumption of positive costs implies that the two strategies of getting informed and then ignoring the information are strictly dominated by the strategy of not getting informed and going to the same food source.

6 For instance 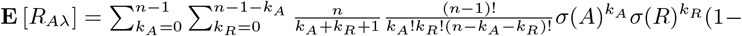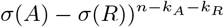

7 In a setting in which individuals can have different degrees of responsiveness one might want to assume that the cost of responsiveness varies with the degree of responsiveness. We shall not pursue this here, however.

8 The distribution with cdf 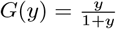 is the distribution of a random variable *Y* such that its reciprocal (or inverse)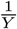 has exactly the same distribution. One could call *G* the inversion invariant distribution.

9 Hofbauer and Sandholm [2009] actually use the letter *n* instead of *k*, but for our paper *n* is reserved to denote the number of players.

